# Integrative Machine Learning and Bioinformatics Analysis to Identify Cellular Senescence-Related Genes and Potential Therapeutic Targets in Ulcerative Colitis and Colorectal Cancer

**DOI:** 10.1101/2025.03.24.644876

**Authors:** Tianle Xue, Yunpeng Chen, Xiaomeng Li, Zhixiang Zhou

## Abstract

**Background:** Ulcerative colitis (UC) is a chronic inflammatory condition that predisposes patients to colorectal cancer (CRC) through mechanisms that remain largely undefined. Given the pivotal role of cellular senescence in both chronic inflammation and tumorigenesis, we integrated machine learning and bioinformatics approaches to identify senescence□related biomarkers and potential therapeutic targets involved in the progression from UC to CRC.

**Methods:** Gene expression profiles from six GEO datasets were analyzed to identify differentially expressed genes (DEGs) using the limma package in R. Weighted gene co expression network analysis (WGCNA) was employed to delineate modules significantly associated with UC and CRC, and the intersection of DEGs, key module genes, and senescence related genes from the CellAge database yielded 112 candidate genes. An integrated machine learning (IML) model, utilizing 12 algorithms with 10 fold cross validation was constructed to pinpoint key diagnostic biomarkers. The diagnostic performance of the candidate genes was evaluated using receiver operating characteristic (ROC) analyses in both training and validation cohorts. In addition, immune cell infiltration, protein protein interaction (PPI) networks, and drug enrichment analyses, including molecular docking were performed to further elucidate the biological functions and therapeutic potentials of the identified genes.

**Results:** Our analysis revealed significant transcriptomic alterations in UC and CRC tissues, with the turquoise module demonstrating the strongest association with disease traits. The IML approach identified five pivotal genes (ABCB1, CXCL1, TACC3, TGFBI, and VDR) that individually exhibited AUC values >0.7, while their combined diagnostic model achieved an AUC of 0.989. Immune infiltration analyses uncovered distinct immune profiles correlating with these biomarkers, and the PPI network confirmed robust interactions among them. Furthermore, drug enrichment and molecular docking studies identified several promising therapeutic candidates targeting these senescence□related genes.

**Conclusions:** This study provides novel insights into the molecular interplay between cellular senescence and the UC to CRC transition. The identified biomarkers not only offer strong diagnostic potential but also represent promising targets for therapeutic intervention, paving the way for improved clinical management of UC associated CRC.

## 1. Introduction

Colorectal cancer (CRC) stands as one of the foremost causes of cancer-related morbidity and mortality globally, posing a significant challenge to public health^[1–3]^. Ulcerative colitis (UC), a chronic inflammatory bowel disease, not only drastically impairs patients’ quality of life but also escalates the risk of developing CRC over time^[4–5]^. Studies have shown that prolonged duration of UC increases the likelihood of CRC occurrence^[6]^, with cell senescence playing a pivotal role in the carcinogenic process^[7]^. However, at the level of cellular senescence, current research on the key genes of colorectal cancer and ulcerative colitis is not clear, and there is currently no study analyzing the relationship between the two diseases and cell senescence from a genomic perspective. Therefore, the research on related genes and the development of drugs are crucial.

Cell Senescence (CS) is a complex biological process in which the gradual decline in physiological functions increases susceptibility to diseases such as cancer^[8–9]^. Genes that induce cellular aging often become overexpressed in human tissues with age, and are significantly overexpressed in anti longevity and tumor suppressor genes, while genes that inhibit cellular aging overlap with longevity promoting genes and oncogenes^[10–11]^. Aging cells release pro-inflammatory cytokines and other factors known as senescence associated secretory phenotype (SASP), which lead to chronic inflammation, impaired tissue regeneration, aging, and age-related diseases, like cancer^[12]^. Understanding the determinants of cellular aging and its correlation with aging is crucial for dissecting the potential mechanisms of aging and age-related diseases, as well as exploring potential therapeutic pathways.

CellAge is a manually curated database that contains 279 human genes that drive cellular aging^[13]^. It was compiled after conducting scientific literature searches on gene manipulation experiments in primary, immortalized, or cancer human cell lines that induce or inhibit CS in cells^[14]^. CellAge aging inducers and inhibitors overlap with oncogenes in the tumor suppressor gene (TSG) database (TSGene 2.0) and ONGene database, and can therefore be used to study cancer-related genes^[15–16]^. By excavating deeply into the databases related to cellular senescence, we can gain a more profound understanding of the relevant processes involved in aging and their roles in diseases.

Machine learning (ML) helps humans learn patterns from complex data to predict future behavioral outcomes and trends^[17]^. ML is widely used for variable filtering and variable selection^[18]^. Previously, research commonly used a single ML algorithm or two integrated ML algorithms (such as artificial neural networks^[19]^, support vector machines^[20]^ and gradient boosting machines^[21]^) to optimize variables. However, a single or only two integrated ML algorithms may miss important potential genes, while integrated ML (IML) methods have more advantages in variable screening and model construction^[22]^. In this study, we focus on studying UC and CRC, using bioinformatics methods combined with IML to investigate in detail the related genes of UC and CRC at the cellular aging level, explore the genetic and transcription factors of UC and CRC, and predict potential therapeutic drugs.

## 2. Methods

### 2.1 Selection of Datasets

Datasets were downloaded from the NCBI Gene Expression Omnibus (GEO; https://www.ncbi.nlm.nih.gov/geo/) using the keywords “Colorectal Cancer” or “Ulcerative Colitis.” Our data analysis process is demonstrated in Figure 1. Detailed information for each dataset, including microarray platform, sample groups, accession numbers, and sample sizes—was recorded. Only datasets containing colon tissue samples from patients with colorectal cancer and ulcerative colitis were included. A total of 6 datasets, namely, GSE52060^[23]^, GSE87211^[24]^, GSE90627^[25]^, GSE36807^[26]^, GSE53306^[27]^, and GSE13367^[28]^, were integrated for this study. The training set were selected as GSE52060, GSE87211, GSE36807, and GSE53306. The testing set were selected as GSE90627 and GSE13367. The details for all datasets are presented in Supplementary Table S.1. To correct for batch effects from different studies, we used the “ComBat” function in the “sva” package (version 3.5.0)^[29–30]^. The effectiveness of batch correction was evaluated by comparing data quality before and after adjustment using principal component analysis (PCA)^[31]^.

**Figure 1.**
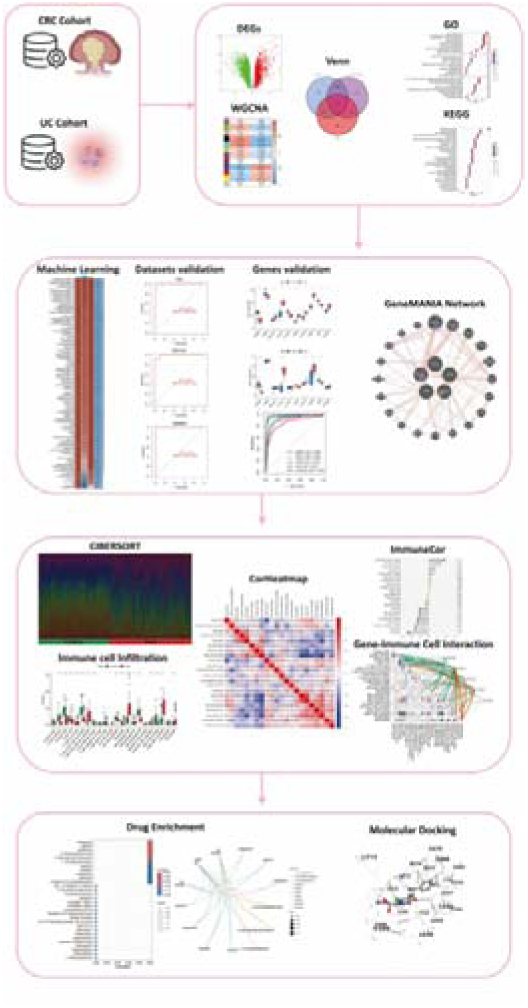
Comprehensive Analysis Workflow for the Study of Colitis-Associated Colorectal Cancer (CRC) Transformation. The workflow includes the analysis of CRC and UC cohorts, identification of differentially expressed genes (DEGs), weighted gene co-expression network analysis (WGCNA), integration through Venn diagrams, and functional enrichment analysis via Gene Ontology (GO) and Kyoto Encyclopedia of Genes and Genomes (KEGG). The study further incorporates machine learning for dataset validation, gene validation using GeneMANIA network analysis, immune cell infiltration assessment through CIBERSORT, and visualization with correlation heatmaps (CorHeatmap). Gene-immune cell interaction is examined using ImmuneCor, followed by drug enrichment analysis and molecular docking to explore therapeutic potentials.

### 2.2 Identification of Differentially Expressed Genes (DEGs) in UC and CRC

To identify key genetic alterations associated with UC and CRC, we performed differential gene expression analysis between case and control groups using the Linear Models for Microarray Data (limma) package in R. Limma is a widely used statistical tool that applies linear models to gene expression data while leveraging empirical Bayes methods to moderate the standard errors of estimated log-fold changes. This approach enhances the stability of statistical inference, particularly in studies with small sample sizes^[32]^. To determine significantly differentially expressed genes (DEGs), we utilized the eBayes function, which computes moderated t-statistics, F-statistics, and log-odds of differential expression for each gene. Genes were considered significantly differentially expressed if they met the threshold of a false discovery rate (FDR) < 0.05 (adjusted p-value < 0.05) and demonstrated an absolute fold change (FC) greater than 0.585 (|log[FC| > 0.5). These stringent criteria helped ensure the robustness and reliability of our findings, highlighting genes with substantial expression changes that may play critical roles in UC and CRC pathogenesis.

### 2.3 Construction of Gene Co-expression Networks Using Weighted Gene Co-expression Network Analysis (WGCNA)

To explore functional gene relationships and identify disease-associated modules, we performed Weighted Gene Co-expression Network Analysis (WGCNA). This method constructs gene co-expression networks and detects modules of highly correlated genes, often linked to specific biological traits^[33]^. As a crucial preprocessing step to ensure a scale-free network topology, we determined the optimal soft-thresholding power (β), selecting a β value where the scale-free topology fit index (R²) exceeded 0.8. A minimum module size of 60 genes was set to identify meaningful gene clusters.

Next, the adjacency matrix was transformed into a Topological Overlap Matrix (TOM), which enhances network robustness by reducing the effects of noise and spurious correlations. To identify gene clusters, we calculated the TOM-based dissimilarity measure (1 - TOM) and applied hierarchical clustering to group genes with similar expression patterns into modules. To refine module detection, dynamic tree cutting was implemented to segment the clustering dendrogram. To assess biological relevance, we correlated module eigengenes (principal components of modules) with clinical traits of UC and CRC. Modules with the strongest correlations and lowest p-values were selected for further analysis, helping identify key gene clusters involved in disease mechanisms and potential therapeutic targets.

### 2.4 Acquisition of senescence related genes in UC and CRC

A comprehensive list of cellular senescence-associated genes was obtained from the CellAge database. By intersecting the gene sets from WGCNA modules, DEGs, and the CellAge dataset via “ggvenn” package (v 0.1.9)^[34]^, we extracted a subset of genes that are not only involved in cellular senescence but also exhibit differential expression and co-expression patterns in UC and CRC. These intersecting genes were considered as potential senescence-related biomarkers and therapeutic targets for further analysis.

### 2.5 Gene set enrichment analysis on functions and pathways

Gene Ontology (GO) provides a structured, dynamically updated vocabulary encompassing gene product attributes across all species, in which GO enrichment contained 3 parts: biological processes (BPs), cellular components (CCs) and molecular functions (MFs)^[35]^. Kyoto Encyclopedia of Genes and Genomes (KEGG) integrates genomic, chemical, and systemic functional information, offering insights into the network of molecular interactions in the cells. For the purpose of understanding candidate genes’ function as well as participating pathways, “clusterProfiler” package (v 4.7.13) was employed for GO and KEGG analysis^[36]^. Utilizing GO and KEGG pathway analyses, we systematically explore the functional and interactive networks that characterize the senescence landscape in UC transitioning into CRC.

### 2.6 Construction and Validation of the Integrated Machine Learning (IML) Model

We developed the final predictive model with optimal performance by applying 10-fold cross-validation on the training set, evaluating 113 model combinations derived from 12 machine learning algorithms. These algorithms included Lasso, Ridge, Stepwise GLM (Stepglm), Random Forest (RF), XGBoost, Elastic Net (Enet), Linear Discriminant Analysis (LDA), Partial Least Squares Regression for Generalized Linear Models (plsRglm), Generalized Boosted Regression Models (GBM), Naive Bayes, GLMBoost, and Support Vector Machine (SVM). The 113 models consisted of 22 individual algorithms and 91 combined algorithms, as detailed in Supplementary Table S.2. To determine the best-performing model, we calculated the concordance index (C-index) for each model and selected the one with the highest C-index as the optimal model. The genes identified by this model were considered candidate disease-related genes, potentially serving as biomarkers for UC and CRC.

After constructing the integrated machine learning (IML) model, we assessed its classification performance using confusion matrices for the training set and two independent validation datasets, GSE13367 and GSE90627. To further validate the model’s predictive capability, we generated Receiver Operating Characteristic (ROC) curves for both the training and validation sets and computed the Area Under the Curve (AUC) with 95% Confidence Intervals (CI). A model was deemed statistically rational only if the ROC AUC exceeded 0.7 for both the training and validation sets^[37–38]^. This approach ensured the robustness and generalizability of our model in distinguishing disease-associated genes and validating their diagnostic potential.

### 2.7 Differential Gene Expression Analysis and ROC Curve Construction

Differential gene expression analysis was performed using experimental data from the GEO datasets. To compare the expression levels of disease-related genes between the UC and CRC validation cohorts, we conducted Student’s t-test. Genes exhibiting statistically significant differential expression (p < 0.05) were identified as cellular senescence-related genes in UC or CRC. To evaluate their diagnostic potential, we generated Receiver Operating Characteristic (ROC) curves for each gene and calculated the Area Under the Curve (AUC) with 95% Confidence Intervals (CI). Genes with an AUC greater than 0.7 in both UC and CRC patients were considered to have significant diagnostic value^[39]^. Furthermore, the significantly differentially expressed genes were integrated into a combined diagnostic model, and its ROC curve was constructed. If the combined model exhibited an AUC higher than that of any individual gene, it was considered a more effective diagnostic tool. The volcano plot was redrawn to visualize the upregulation or downregulation of genes with significant expression differences between UC and CRC.

### 2.8 Construction of the Protein-Protein Interaction (PPI) Network

A protein-protein interaction (PPI) network was constructed to explore the functional relationships and interaction dynamics among the genes with significant expression differences identified in IML. GeneMANIA (http://genemania.org/) incorporates data from multiple interaction types, including co-expression, physical interactions, genetic interactions, co-localization, pathway participation, and shared protein domains, providing a holistic view of the gene interactions. The genes with significant expression differences were input into GeneMANIA to generate a comprehensive PPI network.

### 2.9 Analysis for Immune Cell Infiltration

To investigate disparities in immune infiltration between the two risk groups, the infiltration abundance of 22 distinct immune cell types^[40]^ was first quantified using the CIBERSORT algorithm as implemented in the IOBR package (v 0.99.9)^[41]^. A Wilcoxon test was then applied to identify immune cell populations displaying significant differences (p < 0.05) between the risk groups. Subsequently, Spearman correlation analyses were performed with the psych package (v 2.4.3), using thresholds of |cor| > 0.3 and p < 0.05, to elucidate the correlation network among these differentially abundant immune cells. In addition, correlations between these immune cells and prognostic genes were evaluated under the same thresholds to further characterize the interplay between immune infiltration and gene expression profiles.

### 2.10 Identification of novel drug targets

To explore potential therapeutic agents targeting cellular senescence related genes in UC and CRC, we conducted a drug enrichment analysis using Enrichr (https://maayanlab.cloud/Enrichr/). This analysis, with a significance threshold of p<0.05, aimed to identify compounds that specifically interact with these genes, providing valuable insights for drug repurposing and novel treatment strategies. Furthermore, to evaluate the molecular interactions between candidate drugs and target proteins, we utilized CB-Dock2, an advanced version of the CB-Dock server designed for protein-ligand blind docking. CB-Dock2 integrates cavity detection, molecular docking, and homologous template fitting, offering a more precise and automated approach for structure-based drug discovery. Given the three-dimensional (3D) structures of a protein and a ligand, the platform predicts their binding sites and binding affinity, facilitating computer-aided drug discovery and optimization (https://cadd.labshare.cn/cb-dock2/index.php). By combining drug enrichment analysis with molecular docking, our study enhances the identification of potential therapeutic candidates and accelerates the development of targeted treatments for UC and CRC.

## 3. Results

### 3.1 Acquisition of senescence related genes in UC and CRC

All diseased samples from the training set (GSE52060, GSE87211, GSE36807, and GSE53306) were merged into “Treat”, and all healthy control samples were merged into “Control”. In the heatmap, the validation group and experimental group are divided into different modules, and the samples in each dataset are segmented into different squares (Figure 2A). The colors of the squares represent the changes in gene expression, with red representing upregulation and blue representing downregulation. A total of 3446 DEGs were identified by comparing the BD and control groups. Among all DEGs, 1716 genes displayed upregulation, whereas 1730 genes were downregulated (Figure 2B).

**Figure 2.**
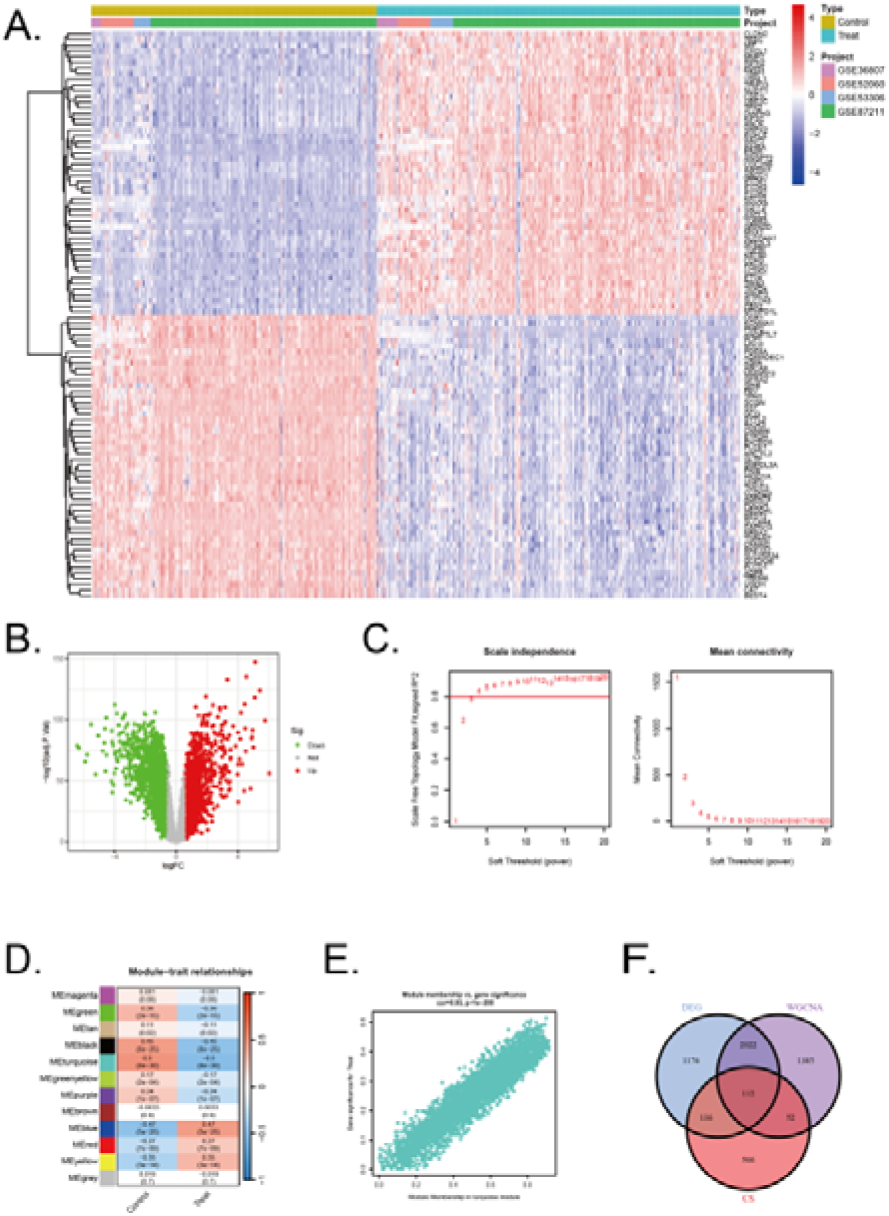
Identification of DEGs in UC/CRC patients and identification of key genes by WGCNA analysis in UC/CRC patients. (A) Heatmap showing up-regulated or down-regulated DEGs in UC/CRC samples compared to normal samples (bule: down-regulated; red: up-regulated). (B) Volcano plot of DEGs between UC/CRC and controls. (C) Analysis of network topology for various soft thresholds (β). (D) Module-trait relationships. (E) Associations between turquoise module membership and gene importance is depicted in a scatter plot. (F) The overlapping regions from key module genes, DEGs, and cellular senescence related genes.

Next, WGCNA was used to identify the significant module genes associated with UC and CRC. We selected the optimal soft-thresholding power (β) to establish a scale-free topology network, ensuring that the scale-free topology fit index (R²) exceeded 0.8 (Figure 2C). The chosen β value was set to maintain the network’s scale-free characteristics. The grey module and brown module did not successfully cluster the genes commonly considered irrelevant or uninformative (i.e., the “junk module”). The turquiose (r[=[0.5, p[=8×10^−30^) module displayed the highest correlation with UC and CRC (Figure 2D). The relationship between module membership and gene significance in the turquiose module is calculated (Cor=0.93, p<10^−200^) and plotted(Figure 2E). Consequently, 3571 significant module genes were identified.

Through a comprehensive analysis integrating data from the CellAge database, weighted gene co-expression network analysis (WGCNA) modules, and differentially expressed genes (DEGs), we identified a refined subset of 112 shared genes implicated in cellular senescence and their association with UC and CRC. The Venn diagram (Figure 2F) summarizes the intersection results, illustrating the overlap between the gene sets and emphasizing the genes that could serve as pivotal links between cellular senescence and disease progression in UC and CRC.

### 3.2 Functional Annotation and Pathway Enrichment Analysis

The differentially expressed genes identified using the limma R package were analyzed through Gene Ontology (GO) enrichment analysis (Figure 3A, 3B). The results were ranked in ascending order based on adjusted p-values (p.adjust) and GeneRatio. In the Biological Processes (BP) category, the top three pathways with the lowest p.adjust values and the highest number of enriched genes were morphogenesis of a branching structure, morphogenesis of a branching epithelium, and gland development. In the cellular components (CC) category, the top 3 terms were cytoplasmic vesicle lumen, secretory granule lumen, and collagen − containing extracellular matrix. In the molecular functions (MF) category, the top 3 terms were DNA− binding transcription factor binding, DNA− binding transcription activator activity, DNA−binding transcription activator activity, RNA polymerase II−specific.

**Figure 3.**
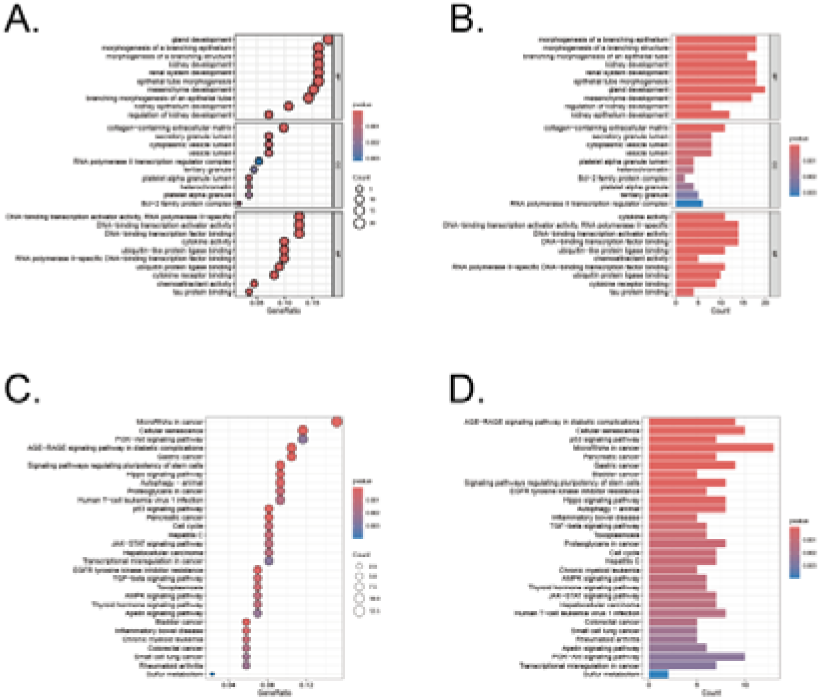
GO and KEGG analysis of the overlapping genes. (A-B) GO analysis of these overlapping genes in UC/CRC patients. (C-D) KEGG analysis of these overlapping genes in UC/CRC patients.

The KEGG pathway analysis revealed key pathways that were significantly enriched among the genes identified in our study (Figure 3C, 3D). These pathways included the PI3K-Akt signaling pathway, p53 signaling pathway, and the cell cycle, which are known to play pivotal roles in regulating cellular senescence, survival, proliferation, and apoptosis. The involvement of these pathways underscores the potential mechanisms through which cellular senescence could influence the transition from UC to CRC.

### 3.3 Identification of Intersection genes with diagnostic value and developing a diagnostic model for UC-related CRC via machine learning

A comprehensive machine learning approach involving 12 algorithms was implemented with a 10-fold cross-validation process to identify the most robust diagnostic model based on 112 shared genes (Figure 4A). The analysis was conducted using the training dataset and validated across two external datasets (GSE90672 and GSE13367). The final model, which demonstrated the best performance, was constructed by integrating Stepglm[both] and Enet[α=0.6]. Specifically, the Stepglm[both] algorithm identified 10 pivotal genes, including ABCB1, AGR2, BCL2L1, CXCL1, FOXO1, SOX4, TACC3, TGFβI, VDR, and VEGFA, while the Enet[α=0.6] algorithm optimized the model’s reliability. The calibration curves, illustrated in Fig. 6D and Fig. 6E, show high AUC values for the training set (Figure 4B, AUC=0.991), as well as the testing set GSE90627 (Figure 4C, AUC=1.000) and GSE13367 (Figure 4D, AUC=0.993), indicating a strong agreement between the predicted probabilities and observed clinical outcomes. These results highlight the robust calibration and diagnostic performance of the proposed model.

**Figure 4.**
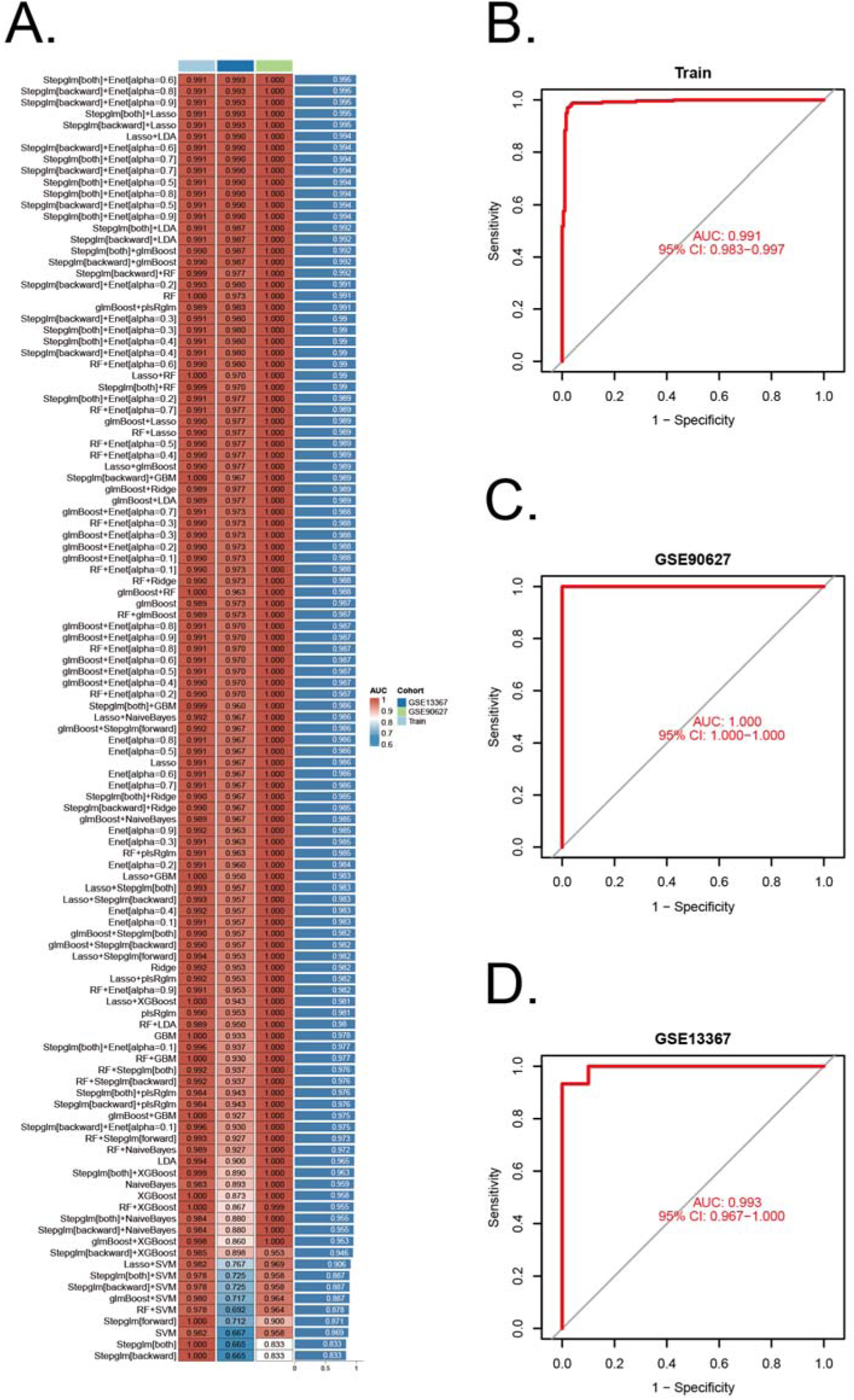
Construction and validation of diagnostic signatures by integrative machine learning. (A) The 113 combinations of prediction models using 10-fold cross-validation with ranked AUC index. (B-D) ROC plots for datasets in internal training set and external validation sets (GSE90672 and GSE13367), correspondingly.

### 3.4 Diagnosis value of pivotal genes

Ten pivotal genes were included in the following ROC analysis. All 10 pivotal genes showed high significance (p<0.001) in CRC and its control group (Figure 5A), while ABCB1, CXCL1, TACC3, TGFβI, VDR displayed high significance (p<0.001) in UC and its control group(Figure 5B). Based on the significant differences in gene expression, ABCB1, CXCL1, TACC3, TGFβI, and VDR were integrated into a combined model. All the 5 genes were included in ROC analysis. We calculated the AUC values for each gene and the combined model separately (Figure 5C). The results showed that the AUC values of all genes were not less than 0.7, and the AUC value of the combined model (AUC=0.989) was higher than that of any individual gene. Therefore, the combined model has greater diagnostic value compared to any individual gene.

**Figure 5.**
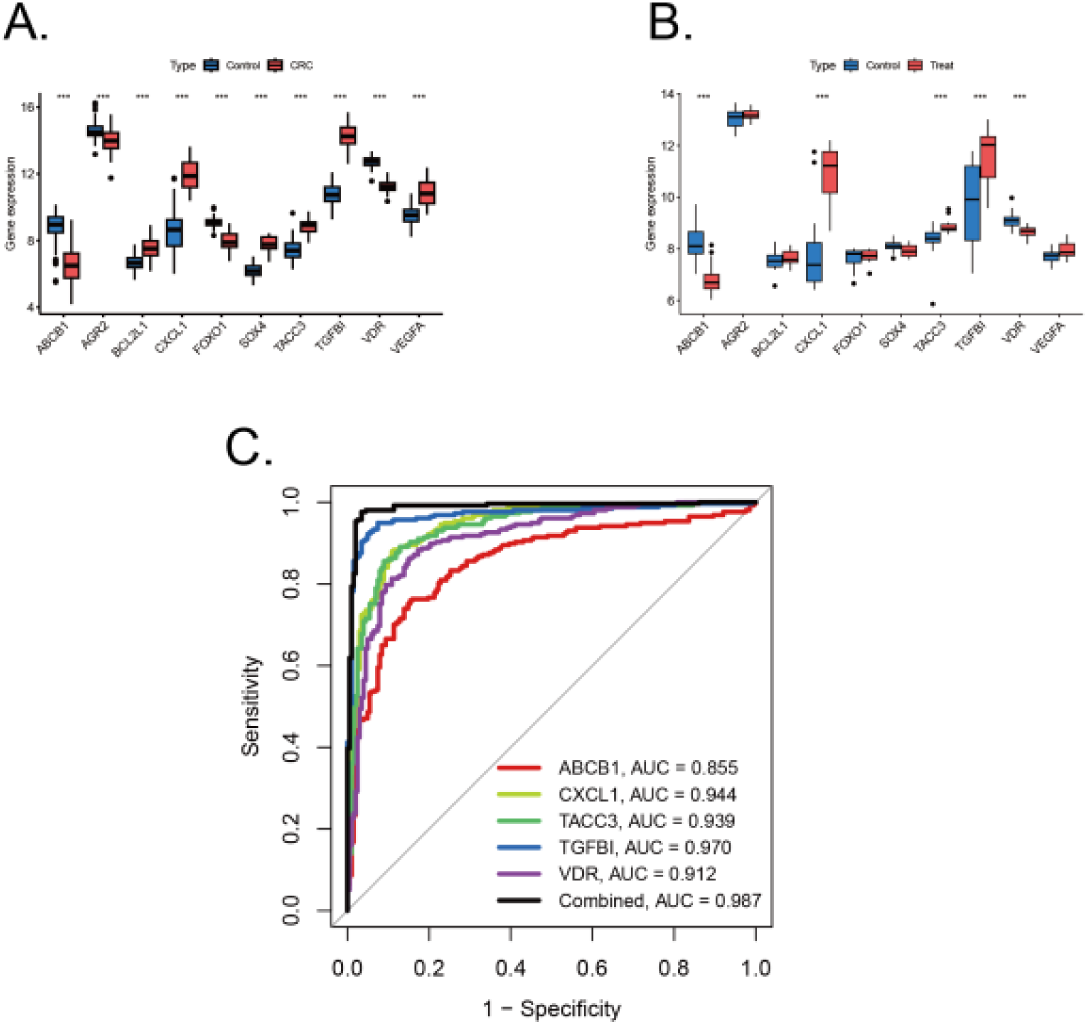
Validation of diagnostic value of pivotal genes. (A) Pivotal genes expression in colorectal cancer training sets with significance (***p<0.001). (B) Pivotal genes expression in ulcerative colitis training sets with significance (***p<0.001). (C) ROC plots for each diagnostic gene and the combined model in internal training cohorts.

**Figure 6.**
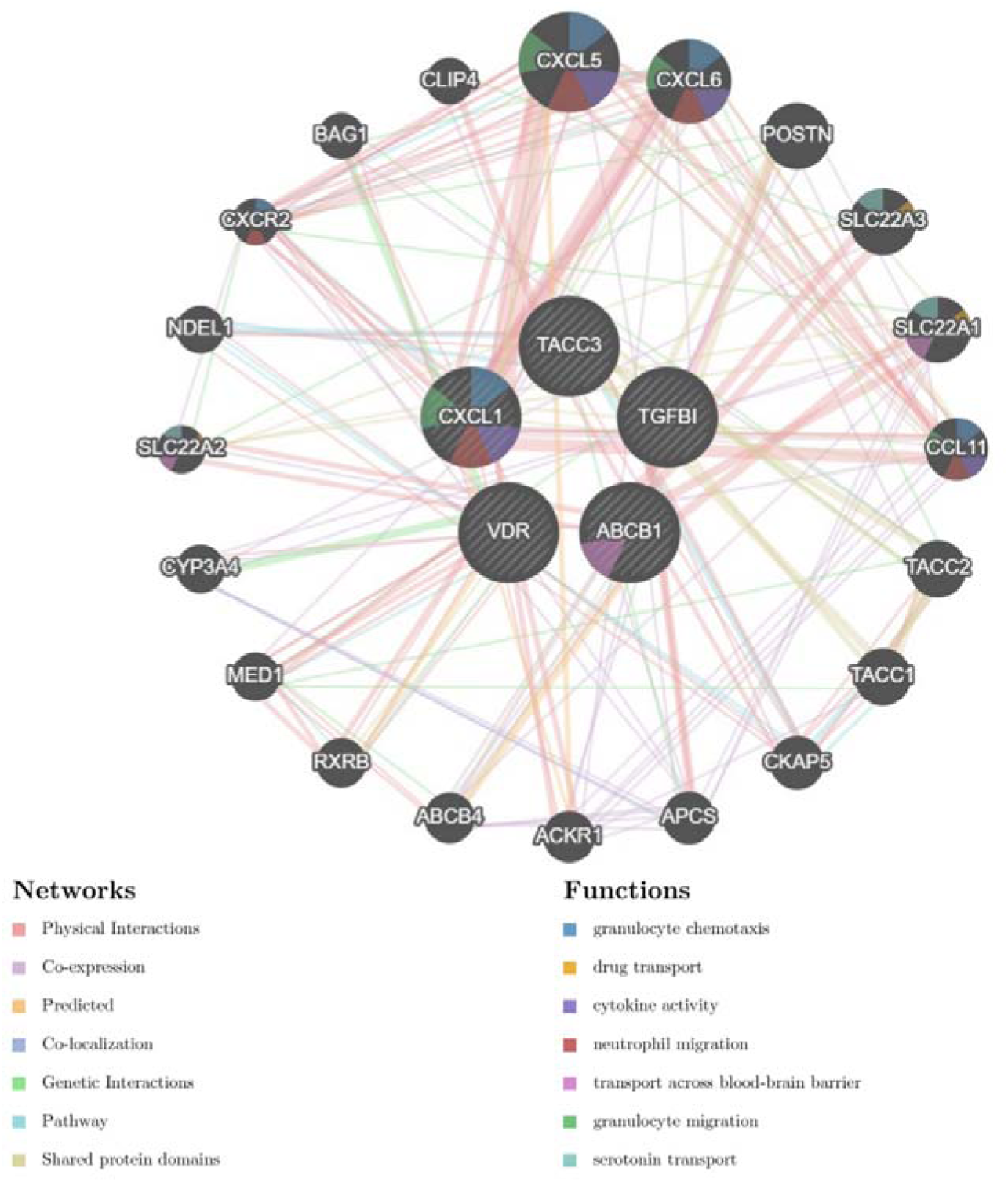
Protein-Protein Interaction (PPI) network for five combined model genes and their related biological functions.

### 3.5 Protein-Protein Interaction (PPI) Network Construction

The PPI network for genes which were included in combined model genes (ABCB1, CXCL1, TACC3, TGFβI, and VDR) were created via the GeneMANIA database (https://genemania.org/). In the GeneMANIA map, a total of 20 genes (CXCL5, CXCL6, POSTN,SLC22A3, SLC22A1, CCL11, TACC2, TACC1, CKAP5, APCS, ACKR1, ACBC4, RXRB, MEDI, CYP3A4, SLC22A2, NDEL1, CSCR2, BAG1, and CLIP4) were found to have gene interactions with five combined model genes (Figure 6). In the GeneMANIA network, physical interactions between pivotal genes and other genes account for 77.64%, while co-expression accounts for 8.01%, demonstrating the strong protein-protein interactions within the GeneMANIA network topology.

### 3.6 Analysis of Immuno-infiltration and Correlation analysis

Immune correlation analysis was performed with all samples in training set (Figure 7A). The infiltration landscape showed that 22 kinds of immune cell distributions in the control and treat groups. Fourteen types of immune cells (neutrophils, mast cells activated, mast cells resting, macrophages M2, macrophages M0, monocytes, NK cells activated, T cells follicular helper, T cells CD4 memory activated, T cells CD4 memory resting, T cells CD4 naive, T cells CD8, and B cells memory) infiltrated significantly (p<0.001) between the control and treat groups (Figure 7B). Correlation analysis between immune cells indicates that Macrophage M2 exhibited significantly negative correlation with activated T cells CD4 naive (r = -0.63, p < 0.05), T cells CD8 had positive correlation with macrophages M2 (r = 0.31, p < 0.05) (Figure 7C). The correlation between genes and 22 immune cell types, as well as the interrelationships among immune cells, has been systematically analyzed and visualized (Figure 7D). Among the findings, ABCB1 exhibits the strongest positive correlation with T cells CD4 memory resting, while demonstrating the most pronounced negative correlation with Monocytes (Figure 7E). Similarly, CXCL1 is most positively correlated with Neutrophils and Macrophages M0, whereas its most significant negative associations are observed with NK cells activated and Macrophages M2 (Figure 7F). In the case of TACC3, its highest positive correlation is identified with Neutrophils, whereas its strongest negative correlations are noted with B cells memory and Plasma cells (Figure 7G). Notably, TGFβI shows the most significant positive correlation with Mast cells activated, while displaying a marked negative correlation with Plasma cells, T cells CD8, T cells gamma delta, and Macrophages M2 (Figure 7H). The gene VDR did not exhibit strong correlations with immune cells in the analysis (Figure 7I).

**Figure 7.**
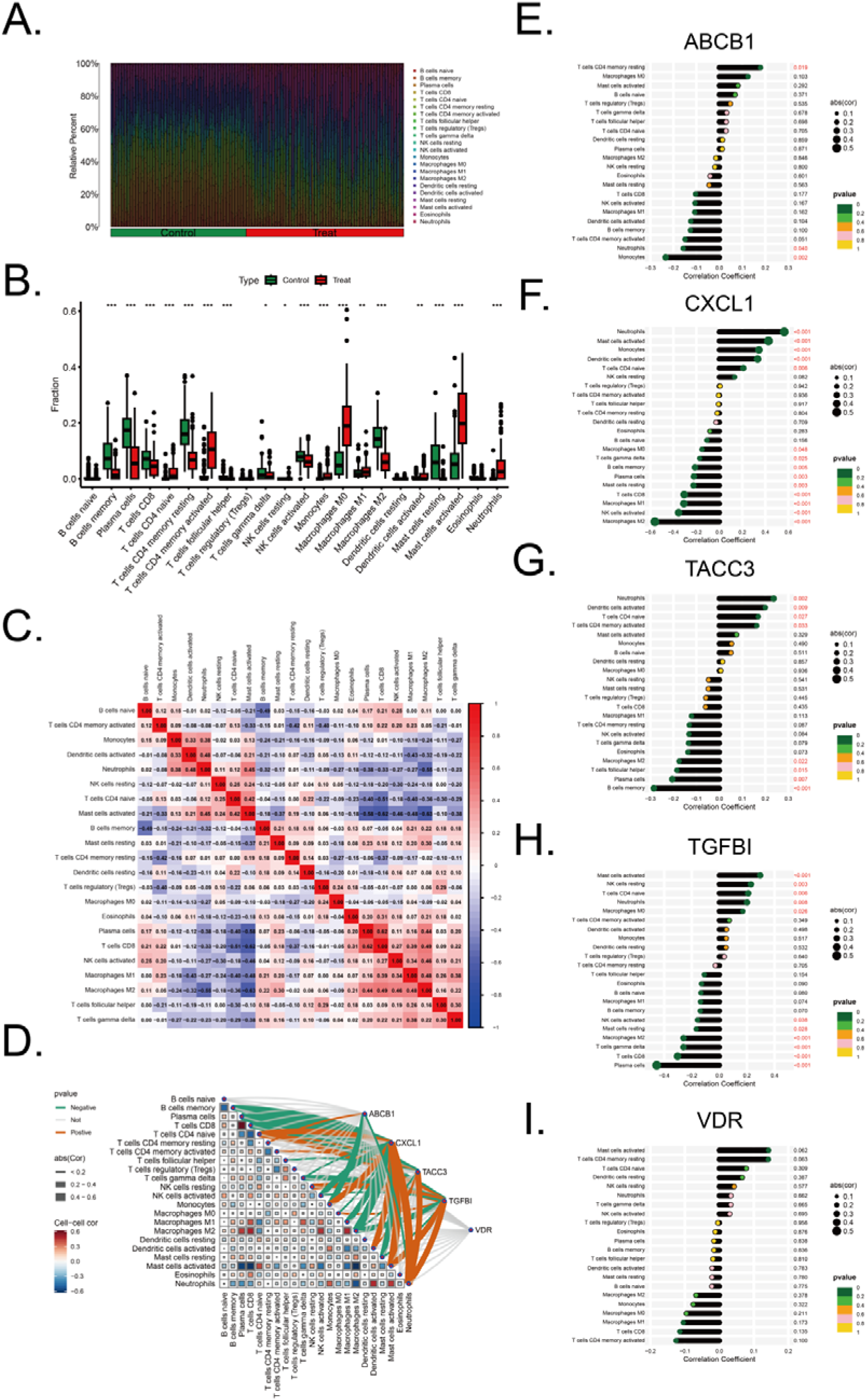
Immune infiltration landscape in colorectal cancer and ulcerative colitis. (A) Proportional graph of 22 kinds immune cells in all training sets. (B) Distribution of different types of immune cells in control group and CRC/UC group (*p<0.05, **p<0.01, ***p<0.001). (C) Correlation of 22 immune cells by compositions. Both horizontal and vertical axes demonstrate immune cells subtypes. (D) Correlation analysis of the level of infiltration of five pivotal genes and each type of immune cells. (E-I) The association between ABCB1, CXCL1, TACC3, TGFβI and VDR expression with different immune cell infiltration in the treat group, correspondingly.

### 3.7 Potential Drug Discovery and Gene-drug Interaction

Enrichr database was utilized to screen therapeutic agents that target the five combined model genes (Table S1). The analysis of the predicted results indicated potential efficacy in targeting the combined model genes associated with UC and CRC. The identified therapeutic agents include iodoquinol, cefaclor, pyrithione, 5-Aminosalicylic acid, 2-Mercaptobenzothiazole, 1,10-Phenanthroline, eugenol, Bisulfite, Alitretinoin, and gossypol (Figure 8A, 8B). Genes CXCL1, TACC3 and VDR are selected as targets, and gene-drug interactions are represented by interconnected curves, the enriched drugs are then connected to their targets to explore the possibility of these drugs as therapeutic measures (Figure 8C). 5-Aminosalicylic acid is already a common drug used in the treatment of UC. According to the enrichment result, the four most significant drugs with the lowest P value were selected (iodoquinol, cefaclor, pyrithione, 5-Aminosalicylic acid) for visual docking with their targets. Software PyMOL was utilized to visualize the best docking results for four most significant drugs with their targets (Figure 8D-8K).

**Figure 8.**
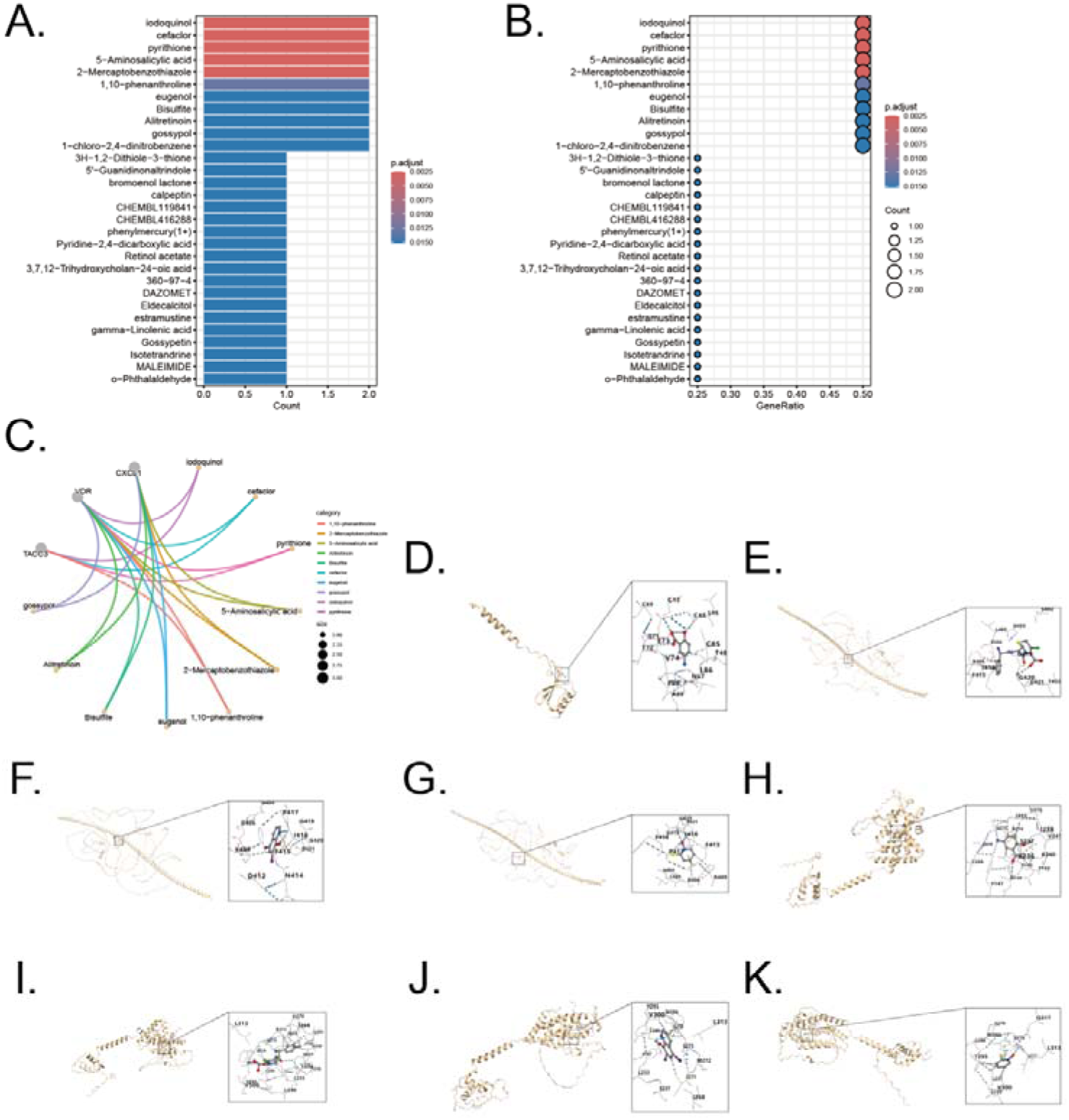
Drug enrichment and molecular docking for five combined model genes. (A-B) Exploration potential drug from Enrichr to five pivotal genes. (C) Visualization of chemical compound data illustrating the distribution and categorization of various pharmaceutical agents. (D) Visualization of molecular docking for 5-Aminosalicylic acid to its target CXCL1. (E) Visualization of molecular docking for cefaclor to its target TACC3. (F) Visualization of molecular docking for iodoquinol to its target TACC3. (G) Visualization of molecular docking for pyrithione to its target TACC3. (H) Visualization of molecular docking for 5-Aminosalicylic acid to its target VDR. (I) Visualization of molecular docking for cefaclor to its target VDR. (J) Visualization of molecular docking for iodoquinol to its target VDR. (K) Visualization of molecular docking for pyrithione to its target VDR.

## 4. Discussion

Sustained ulcerative colitis of the colorectal leads to tissue damage and repair, which is associated with an increased incidence of colitis-associated colorectal cancer. Meanwhile, cellular senescence may be a trigger for colorectal cancer or an emerging therapeutic target^[42]^. To our knowledge, our work is the first to filter senescence-related genes and potential therapeutic drugs in UC and CRC based on the overall normalized weights of IML. Four training sets, two testing set, a total of 621 samples in GEO database were included, and clinical studies were promoted by using datasets to validate the results. Five genes in combined model, ABCB1, CXCL1, TACC3, TGFβI, and VDR, all showed an AUC > 0.7 in gene ROC plot, and their combination diagnostic model showed higher AUC value than any other individual genes, indicating a potential diagnostic value of the five combined model genes. We further investigated the immune correlations of the five genes in the combined model and expanded their potential diagnostic value. These genes highlight the intricate relationship between cellular senescence, immune response, and tumor progression in CRC. Moreover, the development of novel anti-cancer, anti-inflammatory, and anti-aging drugs is often costly and time-consuming. By leveraging bioinformatics to identify medications targeting these key genes, our approach has the potential to enhance efficiency and significantly reduce the costs associated with drug discovery.

ABCB1, also known as P-glycoprotein (P-gp) or MDR1, is a type of ATP-binding cassette (ABC) transporter. The gene encodes a membrane-bound protein that belongs to the ATP-binding cassette (ABC) transporter superfamily. This protein functions as an ATP-driven drug efflux pump, capable of exporting a wide range of xenobiotic compounds due to its broad substrate specificity. ABCB1 helps protect cells from toxic compounds but also contributes to multidrug resistance (MDR) in colorectal cancer cells by reducing the intracellular concentration of anticancer drugs, including doxorubicin, paclitaxel, and vincristine, making them less effective^[43–44]^. In UC, the dysfunction or low activity of ABCB1 leads to the accumulation of harmful bacterial products within the gut epithelium, contributing to chronic inflammation and mucosal damage. This impaired function disrupts the balance of the gut microbiome, exacerbating the inflammatory response and promoting the development of UC symptoms^[45]^. Furthermore, ABCB1 has been identified as a cell senescence related gene, the expression of ABCB1 may increase during aging to enhance the resistance of cells to external toxic substances^[46]^.

CXCL1 is a potent neutrophil chemoattractant that plays a significant role in the immune response. CXCL1’s function in UC is to facilitate the migration and activation of immune cells, thereby exacerbating the inflammatory response in the colon, which is positively correlated with UC severity^[47]^. CXCL1 plays a significant role in CRC by promoting tumor progression through several mechanisms. It is overexpressed in colorectal cancer tissues and contributes to cancer cell proliferation, migration, and invasion. CXCL1 activates the NF-κB pathway, which is crucial for cancer cell survival and inflammation^[48]^. Additionally, CXCL1 recruits myeloid-derived suppressor cells (MDSCs) via the CXCL1-CXCR2 axis, which helps the tumor evade the immune system. CXCL1 is part of the senescence-associated secretory phenotype (SASP), involves its role in the tumor microenvironment, and helps wake up dormant cancer cells, making them more aggressive and prone to recurrence^[49]^.

TACC3, a member of the transforming acidic colied-coil protein family, is found to be overexpressed in colorectal cancer tissues, contributing to increased cell proliferation and cellular senescence. TACC3 regulates various processes during mitosis and interphase. During mitosis, it interacts with proteins like KIFC1 to cluster extra centrosomes, preventing multipolar spindle formation and ensuring proper cell division^[50]^. In interphase, TACC3 interacts with the NuRD complex to suppress tumor suppressor genes, promoting cell cycle progression and survival^[50]^. Targeting TACC3 with inhibitors can induce mitotic catastrophe and G1 phase arrest, leading to cancer cell death, making it a promising therapeutic target for aggressive cancers. Additionally, high TACC3 expression is linked to an immunosuppressive tumor microenvironment and higher tumor mutational burden, suggesting its involvement in tumor progression and immune evasion. Knockdown of TACC3 reduces cell proliferation and senescence, indicating its potential as a therapeutic target^[51]^.

TGFβI, or Trnsforming Growth Factor Beta-Induced protein, is a RGD-containing protein that binds to type I, II and IV collagens, playing a significant role in cancer, particularly in colorectal cancer (CRC)^[52]^. It is involved in promoting angiogenesis, which is the formation of new blood vessels, thereby supporting tumor growth and metastasis. TGFβI expression is regulated by TGFβ signaling pathways, and its presence is associated with increased metastatic potential in CRC cells. TGFβI’s interactions with extracellular matrix proteins and integrins are crucial for its role in cancer, influencing cell adhesion, migration, and chemotherapy resistance^[53]^. TGFβI downstream gene TGF-β1 is a key cytokine involved in the development of kidney diseases and can induce the expression of p21, a protein that can regulate cell cycle arrest and senescence^[54]^.

The Vitamin D receptor (VDR) plays a crucial role in both colorectal cancer (CRC) and ulcerative colitis (UC). In CRC, VDR helps regulate the immune response and inflammation, which are key factors in cancer progression. VDR deficiency is linked to more severe colitis and an increased risk of developing colorectal cancer. It modulates macrophage polarization, promoting an anti-tumor M1 phenotype over the pro-tumor M2 phenotype^[55]^. This regulation helps prevent the transition from chronic colitis to colorectal cancer. In UC, VDR’s role is similar, as it helps control inflammation and maintain intestinal barrier integrity, reducing the risk of cancer development^[56]^. M1 macrophages have anti-tumor functions, which help in reducing inflammation and preventing the progression of colitis-associated colorectal cancer. The absence of VDR accelerates the progression from chronic colitis to colorectal cancer, highlighting its protective role in this transition. VDR plays a crucial role in regulating DNA repair during oncogene-induced senescence (OIS)^[57]^. When VDR levels are reduced, as seen in cells expressing oncogenic Ras, it leads to a decrease in the DNA repair factors BRCA1 and 53BP1. This reduction impairs the cell’s ability to repair DNA damage, contributing to genomic instability. VDR helps maintain the balance of these repair factors, and its downregulation can exacerbate DNA repair deficiencies, promoting senescence and potentially leading to tumorigenesis^[57]^.

## 5. Conclusion

In this work, we have effectively applied integrative machine learning and bioinformatics approaches to identify key cellular senescence-related genes—namely, ABCB1, CXCL1, TACC3, TGFβI, and VDR—that show promising potential as diagnostic biomarkers and therapeutic targets in the progression from ulcerative colitis to colorectal cancer. Our combined diagnostic model, which outperformed individual gene markers, underscores the significant diagnostic value of these candidates, while our immune infiltration analyses further suggest that immunological dysregulation may play a crucial role in disease evolution. However, the current findings are primarily based on retrospective dataset analyses and predictive modeling, and thus additional experimental and clinical validations are required to fully ascertain the clinical applicability of these genes. In the future, we plan to expand our study with larger, diverse clinical cohorts and mechanistic investigations to further elucidate the roles of these senescence-related genes in UC and CRC, ultimately paving the way for more targeted and effective therapeutic strategies.

## Reference

1. Arnold M, Sierra MS, Laversanne M, Soerjomataram I, Jemal A, Bray F. Global patterns and trends in colorectal cancer incidence and mortality. Gut. 2017 Apr;66(4):683–691.

2. GBD 2019 Colorectal Cancer Collaborators. Global, regional, and national burden of colorectal cancer and its risk factors, 1990-2019: a systematic analysis for the Global Burden of Disease Study 2019. Lancet Gastroenterol Hepatol. 2022 Jul;7(7):627–647.

3. Musa M, Ali A. Cancer-associated fibroblasts of colorectal cancer and their markers: updates, challenges and translational outlook. Future Oncol. 2020 Oct;16(29):2329–2344.

4. Shah SC, Itzkowitz SH. Colorectal Cancer in Inflammatory Bowel Disease: Mechanisms and Management. Gastroenterology. 2022 Mar;162(3):715–730.e3.

5. Yashiro M. Ulcerative colitis-associated colorectal cancer. World J Gastroenterol. 2014 Nov 28;20(44):16389–97.

6. Dan WY, Zhou GZ, Peng LH, Pan F. Update and latest advances in mechanisms and management of colitis-associated colorectal cancer. World J Gastrointest Oncol. 2023 Aug 15;15(8):1317–1331.

7. Risques RA, Lai LA, Himmetoglu C, Ebaee A, Li L, Feng Z, Bronner MP, Al-Lahham B, Kowdley KV, Lindor KD, Rabinovitch PS, Brentnall TA. Ulcerative colitis-associated colorectal cancer arises in a field of short telomeres, senescence, and inflammation. Cancer Res. 2011 Mar 1;71(5):1669–79.

8. Kirkland JL, Tchkonia T. Cellular Senescence: A Translational Perspective. EBioMedicine. 2017 Jul;21:21–28.

9. López-Otín C, Pietrocola F, Roiz-Valle D, Galluzzi L, Kroemer G. Meta-hallmarks of aging and cancer. Cell Metab. 2023 Jan 3;35(1):12–35.

10. Aramillo Irizar P, Schäuble S, Esser D, Groth M, Frahm C, Priebe S, Baumgart M, Hartmann N, Marthandan S, Menzel U, Müller J, Schmidt S, Ast V, Caliebe A, König R, Krawczak M, Ristow M, Schuster S, Cellerino A, Diekmann S, Englert C, Hemmerich P, Sühnel J, Guthke R, Witte OW, Platzer M, Ruppin E, Kaleta C. Transcriptomic alterations during ageing reflect the shift from cancer to degenerative diseases in the elderly. Nat Commun. 2018 Jan 30;9(1):327. doi: 10.1038/s41467-017-02395-2. Erratum in: Nat Commun. 2019 May 31;10(1):2459.

11. Schmitt CA, Wang B, Demaria M. Senescence and cancer - role and therapeutic opportunities. Nat Rev Clin Oncol. 2022 Oct;19(10):619–636.

12. Ou HL, Hoffmann R, González-López C, Doherty GJ, Korkola JE, Muñoz-Espín D. Cellular senescence in cancer: from mechanisms to detection. Mol Oncol. 2021 Oct;15(10):2634–2671.

13. Avelar RA, Ortega JG, Tacutu R, Tyler EJ, Bennett D, Binetti P, Budovsky A, Chatsirisupachai K, Johnson E, Murray A, Shields S, Tejada-Martinez D, Thornton D, Fraifeld VE, Bishop CL, de Magalhães JP. A multidimensional systems biology analysis of cellular senescence in aging and disease. Genome Biol. 2020 Apr 7;21(1):91.

14. Chatsirisupachai K, Palmer D, Ferreira S, de Magalhães JP. A human tissue-specific transcriptomic analysis reveals a complex relationship between aging, cancer, and cellular senescence. Aging Cell. 2019 Dec;18(6):e13041.

15. Zhao M, Kim P, Mitra R, Zhao J, Zhao Z. TSGene 2.0: an updated literature-based knowledgebase for tumor suppressor genes. Nucleic Acids Res. 2016 Jan 4;44(D1):D1023–31.

16. Liu Y, Sun J, Zhao M. ONGene: A literature-based database for human oncogenes. J Genet Genomics. 2017 Feb 20;44(2):119–121.

17. Haug CJ, Drazen JM. Artificial Intelligence and Machine Learning in Clinical Medicine, 2023. N Engl J Med. 2023 Mar 30;388(13):1201–1208.

18. Cascianelli S, Galzerano A, Masseroli M. Supervised Relevance-Redundancy assessments for feature selection in omics-based classification scenarios. J Biomed Inform. 2023 Aug;144:104457.

19. Eetemadi A, Tagkopoulos I. Genetic Neural Networks: an artificial neural network architecture for capturing gene expression relationships. Bioinformatics. 2019 Jul 1;35(13):2226–2234.

20. Huang S, Cai N, Pacheco PP, Narrandes S, Wang Y, Xu W. Applications of Support Vector Machine (SVM) Learning in Cancer Genomics. Cancer Genomics Proteomics. 2018 Jan-Feb;15(1):41–51.

21. Du Z, Zhong X, Wang F, Uversky VN. Inference of gene regulatory networks based on the Light Gradient Boosting Machine. Comput Biol Chem. 2022 Dec;101:107769.

22. Zhang L, Liu Y, Wang K, Ou X, Zhou J, Zhang H, Huang M, Du Z, Qiang S. Integration of machine learning to identify diagnostic genes in leukocytes for acute myocardial infarction patients. J Transl Med. 2023 Oct 27;21(1):761.

23. Medico E, Isella C, Boccaccio C. Expression of the MET oncogene correlates with upregulation of coagulation factor XII and procoagulant disorders in colorectal cancer. 2013; [Internet]. Available from: https://www.ncbi.nlm.nih.gov/geo/query/acc.cgi?acc=GSE52060.

24. Hu Y, Gaedcke J, Emons G, Beissbarth T, Grade M, Jo P, Yeager M, Chanock SJ, Wolff H, Camps J, Ghadimi BM, Ried T. Colorectal cancer susceptibility loci as predictive markers of rectal cancer prognosis after surgery. Genes Chromosomes Cancer. 2018 Mar;57(3):140–149.

25. Guo H, Zeng W, Feng L, Yu X, Li P, Zhang K, Zhou Z, Cheng S. Integrated transcriptomic analysis of distance-related field cancerization in rectal cancer patients. Oncotarget. 2017 May 15;8(37):61107–61117.

26. Montero-Meléndez T, Llor X, García-Planella E, Perretti M, Suárez A. Identification of novel predictor classifiers for inflammatory bowel disease by gene expression profiling. PLoS One. 2013 Oct 14;8(10):e76235.

27. Zhao X, Fan J, Zhi F, Li A, Li C, Berger AE, Boorgula MP, Barkataki S, Courneya JP, Chen Y, Barnes KC, Cheadle C. Mobilization of epithelial mesenchymal transition genes distinguishes active from inactive lesional tissue in patients with ulcerative colitis. Hum Mol Genet. 2015 Aug 15;24(16):4615–24.

28. Bjerrum JT, Hansen M, Olsen J, Nielsen OH. Genome-wide gene expression analysis of mucosal colonic biopsies and isolated colonocytes suggests a continuous inflammatory state in the lamina propria of patients with quiescent ulcerative colitis. Inflamm Bowel Dis. 2010 Jun;16(6):999–1007.

29. Johnson WE, Li C, Rabinovic A. Adjusting batch effects in microarray expression data using empirical Bayes methods. Biostatistics. 2007 Jan;8(1):118–27.

30. Leek JT, Johnson WE, Parker HS, Jaffe AE, Storey JD. The sva package for removing batch effects and other unwanted variation in high-throughput experiments. Bioinformatics. 2012 Mar 15;28(6):882–3. doi: 10.1093/bioinformatics/bts034. Epub 2012 Jan 17.

31. Jolliffe IT, Cadima J. Principal component analysis: a review and recent developments. Philos Trans A Math Phys Eng Sci. 2016 Apr 13;374(2065):20150202.

32. Ritchie ME, Phipson B, Wu D, Hu Y, Law CW, Shi W, Smyth GK. limma powers differential expression analyses for RNA-sequencing and microarray studies. Nucleic Acids Res. 2015 Apr 20;43(7):e47.

33. Langfelder P, Horvath S. WGCNA: an R package for weighted correlation network analysis. BMC Bioinformatics. 2008 Dec 29;9:559.

34. Gao CH, Chen C, Akyol T, Dusa A, Yu G, Cao B, Cai P. ggVennDiagram: Intuitive Venn diagram software extended. Imeta. 2024 Feb 14;3(1):e177.

35. Zhao Y, Huang T, Huang P. Integrated Analysis of Tumor Mutation Burden and Immune Infiltrates in Hepatocellular Carcinoma. Diagnostics (Basel). 2022 Aug 8;12(8):1918.

36. Wu T, Hu E, Xu S, Chen M, Guo P, Dai Z, Feng T, Zhou L, Tang W, Zhan L, Fu X, Liu S, Bo X, Yu G. clusterProfiler 4.0: A universal enrichment tool for interpreting omics data. Innovation (Camb). 2021 Jul 1;2(3):100141.

37. Qin H, Abulaiti A, Maimaiti A, Abulaiti Z, Fan G, Aili Y, Ji W, Wang Z, Wang Y. Integrated machine learning survival framework develops a prognostic model based on inter-crosstalk definition of mitochondrial function and cell death patterns in a large multicenter cohort for lower-grade glioma. J Transl Med. 2023 Sep 2;21(1):588.

38. Chen B, Sun X, Huang H, Feng C, Chen W, Wu D. An integrated machine learning framework for developing and validating a diagnostic model of major depressive disorder based on interstitial cystitis-related genes. J Affect Disord. 2024 Aug 15;359:22–32.

39. Liu C, He Y, Luo J. Application of Chest CT Imaging Feature Model in Distinguishing Squamous Cell Carcinoma and Adenocarcinoma of the Lung. Cancer Manag Res. 2024 Jun 4;16:547–557.

40. Chen F, Yang Y, Zhao Y, Pei L, Yan H. Immune Infiltration Profiling in Nonsmall Cell Lung Cancer and Their Clinical Significance: Study Based on Gene Expression Measurements. DNA Cell Biol. 2019 Nov;38(11):1387–1401.

41. Zhang L, Zhang X, Guan M, Zeng J, Yu F, Lai F. Machine-learning developed an iron, copper, and sulfur-metabolism associated signature predicts lung adenocarcinoma prognosis and therapy response. Respir Res. 2024 May 14;25(1):206.

42. Wu Y, Xie M, Sun JH, Li CC, Dong GH, Zhang QS, Cui PL. Cellular senescence: a promising therapeutic target in colorectal cancer. Future Oncol. 2022 Sep;18(30):3463–3470.

43. Tan P, Xu M, Nie J, Qin J, Liu X, Sun H, Wang S, Pan Y. LncRNA SNHG16 promotes colorectal cancer proliferation by regulating ABCB1 expression through sponging miR-214-3p. J Biomed Res. 2022 Jun 28;36(4):231–241.

44. Lei ZN, Albadari N, Teng QX, Rahman H, Wang JQ, Wu Z, Ma D, Ambudkar SV, Wurpel JND, Pan Y, Li W, Chen ZS. ABCB1-dependent collateral sensitivity of multidrug-resistant colorectal cancer cells to the survivin inhibitor MX106-4C. Drug Resist Updat. 2024 Mar;73:101065.

45. Stoeltje L, Luc JK, Haddad T, Schrankel CS. The roles of ABCB1/P-glycoprotein drug transporters in regulating gut microbes and inflammation: insights from animal models, old and new. Philos Trans R Soc Lond B Biol Sci. 2024 May 6;379(1901):20230074. doi: 10.1098/rstb.2023.0074. Epub 2024 Mar 18.

46. Wu J, Li Y, Nabi G, Huang X, Zhang X, Wang Y, Huang L. Exosome and lipid metabolism-related genes in pancreatic adenocarcinoma: a prognosis analysis. Aging (Albany NY). 2023 Oct 18;15(20):11331–11368.

47. Huo A, Wang F. Biomarkers of ulcerative colitis disease activity CXCL1, CYP2R1, LPCAT1, and NEU4 and their relationship to immune infiltrates. Sci Rep. 2023 Jul 26;13(1):12126.

48. Zhuo C, Ruan Q, Zhao X, Shen Y, Lin R. CXCL1 promotes colon cancer progression through activation of NF-κB/P300 signaling pathway. Biol Direct. 2022 Nov 25;17(1):34.

49. Korbecki J, Gąssowska-Dobrowolska M, Wójcik J, Szatkowska I, Barczak K, Chlubek M, Baranowska-Bosiacka I. The Importance of CXCL1 in Physiology and Noncancerous Diseases of Bone, Bone Marrow, Muscle and the Nervous System. Int J Mol Sci. 2022 Apr 11;23(8):4205.

50. Saatci O, Akbulut O, Cetin M, Sikirzhytski V, Uner M, Lengerli D, O’Quinn EC, Romeo MJ, Caliskan B, Banoglu E, Aksoy S, Uner A, Sahin O. Targeting TACC3 represents a novel vulnerability in highly aggressive breast cancers with centrosome amplification. Cell Death Differ. 2023 May;30(5):1305–1319.

51. Du Y, Liu L, Wang C, Kuang B, Yan S, Zhou A, Wen C, Chen J, Wu Y, Yang X, Feng G, Liu B, Iwamoto A, Zeng M, Wang J, Zhang X, Liu H. TACC3 promotes colorectal cancer tumourigenesis and correlates with poor prognosis. Oncotarget. 2016 Jul 5;7(27):41885–41897.

52. Chiavarina B, Costanza B, Ronca R, Blomme A, Rezzola S, Chiodelli P, Giguelay A, Belthier G, Doumont G, Van Simaeys G, Lacroix S, Yokobori T, Erkhem-Ochir B, Balaguer P, Cavailles V, Fabbrizio E, Di Valentin E, Gofflot S, Detry O, Jerusalem G, Goldman S, Delvenne P, Bellahcène A, Pannequin J, Castronovo V, Turtoi A. Metastatic colorectal cancer cells maintain the TGFβ program and use TGFBI to fuel angiogenesis. Theranostics. 2021 Jan 1;11(4):1626–1640.

53. Corona A, Blobe GC. The role of the extracellular matrix protein TGFBI in cancer. Cell Signal. 2021 Aug;84:110028.

54. Ueda S, Tominaga T, Ochi A, Sakurai A, Nishimura K, Shibata E, Wakino S, Tamaki M, Nagai K. TGF-β1 is involved in senescence-related pathways in glomerular endothelial cells via p16 translocation and p21 induction. Sci Rep. 2021 Nov 4;11(1):21643.

55. Hu PS, Li T, Lin JF, Qiu MZ, Wang DS, Liu ZX, Chen ZH, Yang LP, Zhang XL, Zhao Q, Chen YX, Lu YX, Wu QN, Pu HY, Zeng ZL, Xie D, Ju HQ, Luo HY, Xu RH. VDR-SOX2 signaling promotes colorectal cancer stemness and malignancy in an acidic microenvironment. Signal Transduct Target Ther. 2020 Sep 9;5(1):183.

56. Shi Y, Liu Z, Cui X, Zhao Q, Liu T. Intestinal vitamin D receptor knockout protects from oxazolone-induced colitis. Cell Death Dis. 2020 Jun 15;11(6):461.

57. Graziano S, Johnston R, Deng O, Zhang J, Gonzalo S. Vitamin D/vitamin D receptor axis regulates DNA repair during oncogene-induced senescence. Oncogene. 2016 Oct 13;35(41):5362–5376.

